# Cell-autonomous thermogenesis of macrophage alters its antibacterial function

**DOI:** 10.64898/2026.01.13.699193

**Authors:** Hiroki Sugimoto, Takayuki Isagawa, Kazuhiko Miyanaga, Kotaro Kiga, Yuki Sugiura, Masamichi Yamamoto, Ichiro Manabe, Takahiro Kuchimaru, Longzhu Cui, Norihiko Takeda

**Author notes:** For correspondence: Hiroki Sugimoto, and Norihiko Takeda.

## Abstract

Dynamic temperature gradients exist across the bodies of endothermic animals, from the core to peripheral organ, resulting in the physiological cold environment in superficial regions. Consequently, macrophages distributed throughout the body must be able to adapt not only to thermoneutral conditions but also to colder environments. In fact, it is known that environmental temperature influences macrophage immune responses. However, the thermo-responsive mechanisms of macrophage have been largely unexplored. Here we show that macrophage themselves maintains intracellular temperature under physiological cold condition by increasing proton leak index (defined as mitochondrial proton leak per spare respiratory capacity). We further identified a contribution of ADP/ATP carrier (AAC) to this increase in proton leak index. This cell-autonomous thermogenesis pathway, which does not depend on neural or hormonal inputs, highlights the potential for local and organ-specific temperature regulation. Moreover, cold stress reduced mitochondrial membrane potential, which in turn suppressed the expression of the antimicrobial peptide Resistin-like molecule alpha (RETNLA) and diminished antibacterial properties. Together, these findings suggest that macrophages generate heat whereas compromising antibacterial properties, thereby increasing susceptibility to bacterial infection in physiological cold environment. This adaptation mechanism may underscore the important role of temperature homeostasis in non-adipocyte cells.

## Introduction

Endothermic animals tightly regulate their core body temperature, maintaining it at approximately 37°C. When exposed to low temperatures, they primarily generate heat through muscle shivering (Shivering Thermogenesis) and/or proton leak via uncoupling of mitochondria in brown adipocyte (Non-shivering Thermogenesis) controlled by the sympathetic nervous system (Engel et al, 1992). As a consequence, dynamic temperature gradients arise across the body, ranging from thermoneutral environment at the core to physiological cold condition in peripheral or superficial regions. These peripheral and superficial tissues are particularly susceptible to fluctuations in ambient temperature. For example, the subcutaneous tissues in the human calf and triceps typically maintain a temperature of around 32-33°C but can decrease to around 28°C in cold environments or increase to as high as 37°C in hot conditions or during physical activity (Bazett 1927, Webb 1992). Such ambient temperature changes are thought to directly affect the intracellular temperature and biological functions of macrophages distributed throughout the body in endotherms as well as ectotherms. Therefore, macrophages must be capable of adopting across a wider temperature range. Indeed, environmental temperature has been reported to modulate macrophage immune responses (Salman et. al., 2000, Kizaki et. al., 2002, Lu et. al., 2022). However, the mechanisms by which individual macrophages respond to temperature fluctuations remain poorly understood.

## Results and Discussion

Firstly, to investigate the cold sensitivity of bone marrow-derived macrophages (BMDMs), we conducted a comprehensive metabolomic analysis under physiological cold conditions such as 28°C brought by cold ambient temperature rather than markedly sub-physiological condition such as 4°C. Metabolite profiling revealed a global decrease in metabolite abundance at 28°C compared to 37°C (Fig. 1a). Supervised partial least squares discriminant analysis (PLS-DA) distinguished the metabolomic profiles of cells incubated at 28°C and 37°C, with the primary component separating samples based on temperature (Fig. 1b). Most glycolytic metabolites and fumarate and malate from Tricarboxylic acid cycle were reduced at 28°C compared with 37°C (Fig. 1c). These findings suggest that cold exposure suppresses glycolytic flux and limits electron supply to the electron transport chain of mitochondria. Together, these results support the notion that subtle temperature gradient drastically induces coordinated shifts in both metabolic and mitochondrial processes. To validate these findings at the functional level, we monitored intracellular ATP concentrations using GO-ATeam (Adenosin 5’-Triphosohate indicator based on Epsilon subunit for Analytical Measurements), a genetically encoded ATP biosensor based on green fluorescent protein (GFP) and orange fluorescent protein (OFP) fluorescence resonance energy transfer (Imamura et al., 2009, Nakano et al., 2011). BMDMs from GO-ATeam-expressing mice (AVID mouse, Ohnishi et. al., 2025) were observed by confocal microscopy during temperature shift experiments. Following a temperature shift from 37°C to 28°C, the GO-ATeam fluorescence ratio gradually decreased, indicating a reduction in intracellular ATP levels (Fig. 1d). This observation was consistent with the metabolomic data, confirming that physiological cold exposure leads to global metabolic suppression in BMDMs.

**Fig. 1.**
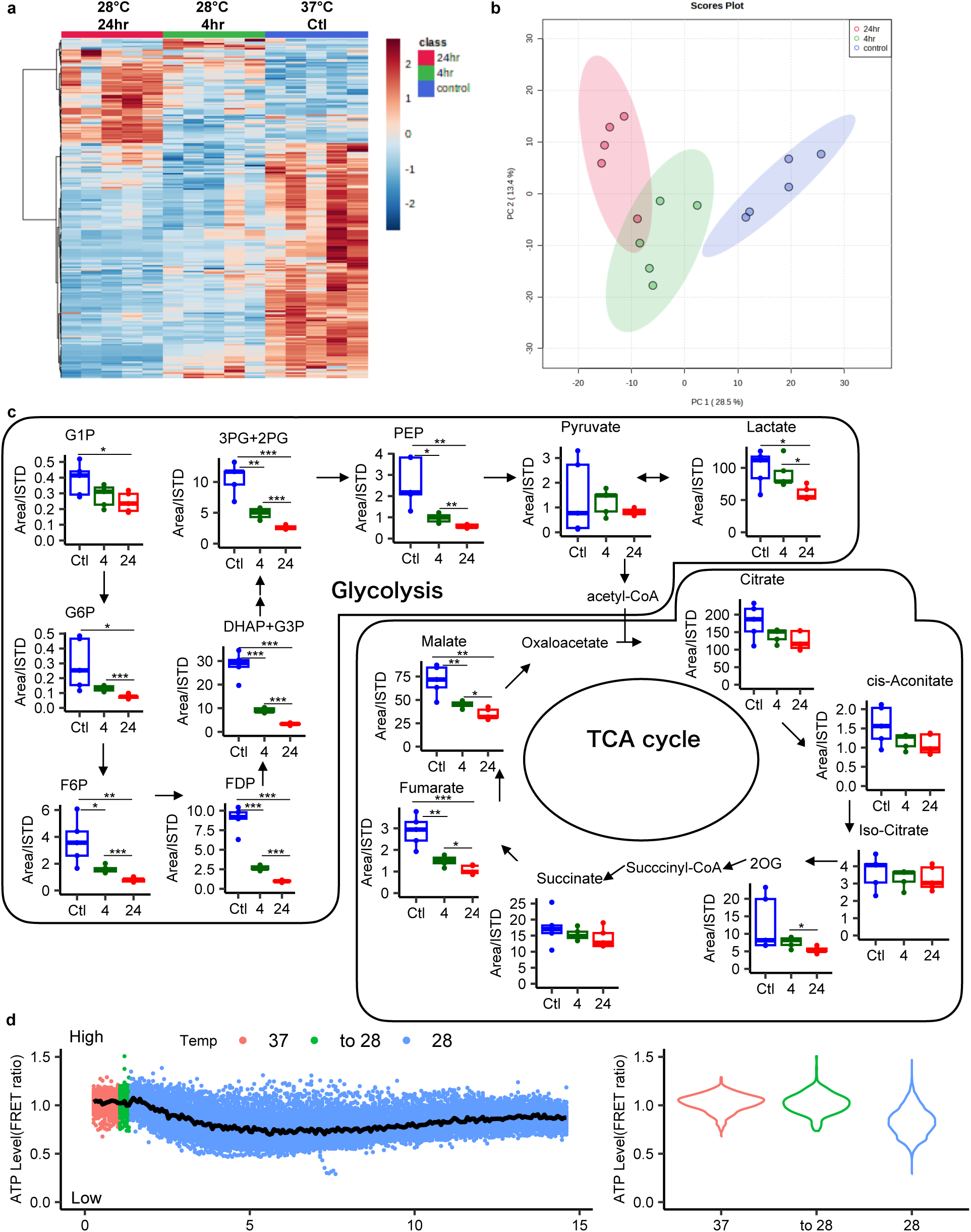
Macrophage responded to cold stimulation. **a**, Heat map of metabolites from BMDMs cultured for 4 and 24 hours in 28°C and in 37°C. n = 5 biological replicates/ group. **b**, Supervised partial least squares discriminant analysis of the metabolites. **c**, Metabolites of glycolysis and TCA cycle compared among 37°C(Ctl, blue), 28°C 4 hours (4, green), and 28°C 24 hours (24,red). n=5 biological replicates/ group. In **c**, box and whisker plots show the the 75th 50th, and 25th percentiles of the data, and minimum and maximum values. **d**, Imaging of ATP biosensor (GO-ATeam) expressing BMDMs. Culture temperature was shifted to 28°C (blue) from 37°C (red). Transition to 28°C (Green). Relative ATP level at start point was set at 1. Abbreviations in **c,** G1P: Glucose-1-phosphate, G6P: Glucose-6-phosphate, F6P: Fructose 6-phosphate, FDP: Fructose1,6-diphosphate, DHAP: Dihydroxyacetone phosphate, G3P: Glyceraldehyde 3-phosphate, PEP: Phosphoenolpyruvate, 3PG: Glycerate 3-phosphat, 2PG: Glycerate 2-phosphat,2OG: 2-oxoglutaric acid.

To explore the impact of global metabolic suppression on mitochondrial function, we measured oxygen consumption rate (OCR) and extracellular acidification rate (ECAR) in BMDMs and four additional cell types: epithelial cells (NCTC1469), keratinocytes (PHK16-0b), white adipocytes (3T3-L1), and brown adipocytes (T37i). OCR and ECAR were used as indicators of mitochondrial respiration and glycolytic activity, respectively. Under physiological cold condition (28°C), all cell types exhibited a reduction in OCR compared to 37°C, indicating a general suppression of cellular metabolism (sFig. 1, Fig. 2a left). Similarly, ECAR was also reduced at 28°C, suggesting that glycolytic activity is likewise inhibited (Fig. 2a middle). Among these, BMDMs appeared to adopt a more quiescent metabolic state in response to cold exposure (Fig. 2a right). In BMDMs, detailed mitochondrial profiling revealed that cold exposure significantly altered several key parameters of mitochondrial respiration—including basal respiration, ATP-linked respiration, maximal respiration, spare respiratory capacity, and proton leak—when comparing 37°C to both 31°C and 28°C (Fig. 2b). Notably, other cell types such as keratinocytes and brown adipocytes also showed decreased proton leak at 28°C compared to 37°C (sFig. 1b, c). Biological reaction rates typically decline by 2–3 times with every 10°C drop in temperature (Hegarty. 1973). This trend was reflected in the basal OCR ratios (37°C/28°C, mean ± standard deviation) across cell types: 1.8□±□0.098 in BMDMs, 2.02□±□0.507 in epithelial cells, 1.54□±□0.335 in keratinocytes, 3.04□±□0.794 in white adipocytes, and 2.23□±□0.660 in brown adipocytes. These data suggest that cold-induced reductions in mitochondrial function are consistent with expected temperature-dependent decreases in biological activity. Interestingly, only BMDMs showed an increased ratio of proton leak to spare respiratory capacity (named as proton leak index) under 28°C conditions compared with 37°C (Fig. 2b), indicating that BMDMs maintain proton leak while experiencing a decline in spare respiratory capacity. Because sex difference of proton leak index in mice was not observed (sFig 2), we used only male mice in the following experiment. Given that proton leak is associated with uncoupling thermogenesis (Rajagopal et al., 2019), its preservation in BMDMs at 28°C suggests enhanced thermogenic activity at lower temperatures. Proton leak serves as an indicator of thermogenic capacity, whereas spare respiratory capacity is considered a marker of mitochondrial health and membrane potential (Windt et al., 2012, Marchetti et al., 2020). Indeed, BMDMs exposed to 28°C exhibited decreased mitochondrial membrane potential, using MT-1, a fluorescence dye sensitive to membrane potential (Fig. 2c). These findings imply that, under physiological cold conditions, BMDMs prioritize thermogenesis over mitochondrial fitness. The regulation of intracellular temperature during ambient temperature changes may be important to maintain the homeostasis in BMDMs.

**Fig. 2.**
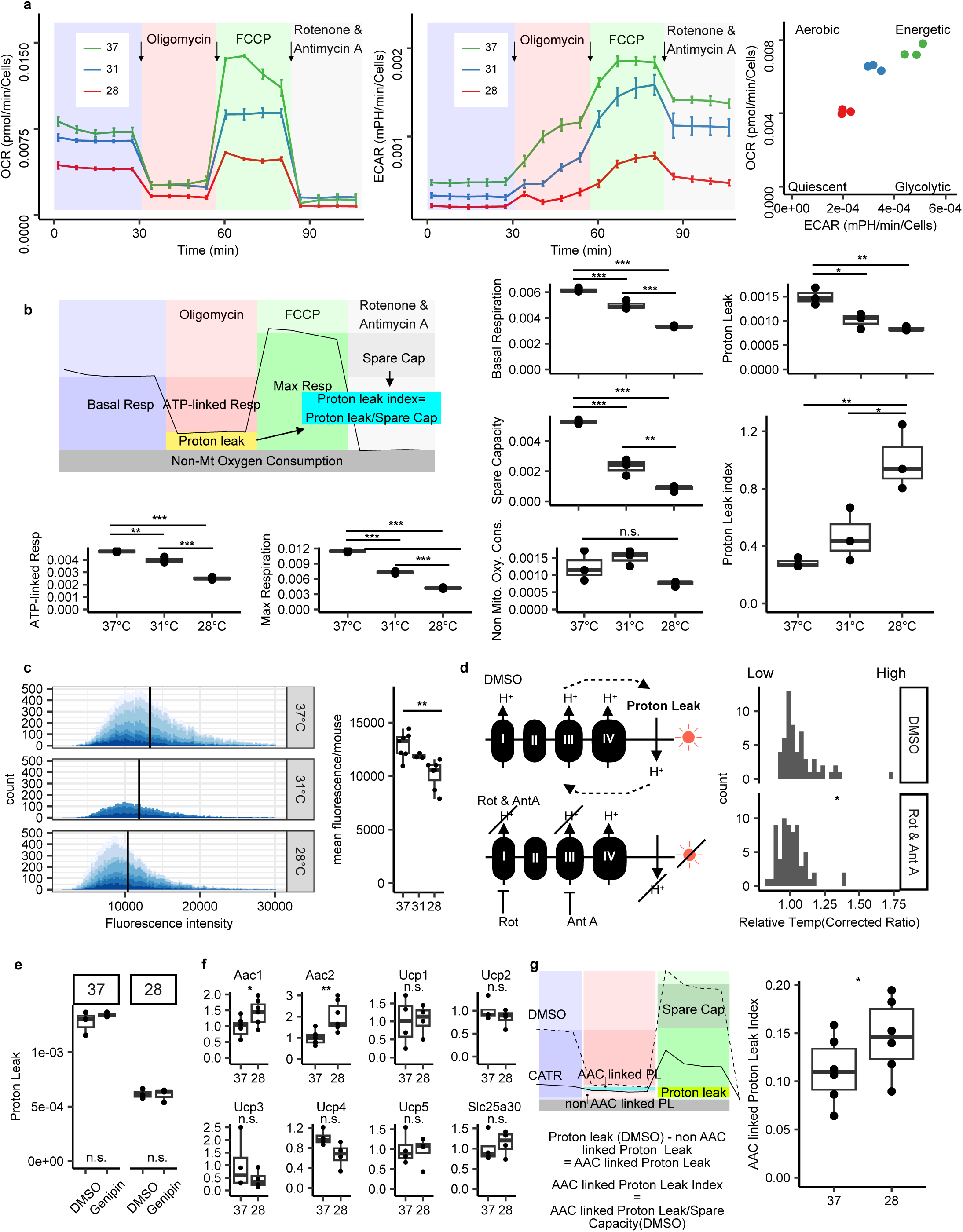
Cold stimulation enhanced proton leak index of macrophage mitochondria. **a**, Respiratory profile of BMDMs (left; OCR, middle; ECAR, right; energy map, n=3 biological replicates/group). **b**, quantification plot from each OCR of BMDMs incubated at 28°C, 31°C, and 37°C(n=3 biological replicates/ group). **c**, left; Histogram of fluorescence intensity of cytoplasmic region per cell, measuring using MT-1 at 28°C, 31°C, and 37°C. Vertical line indicates mean values. Colors indicate biological replicates (n = 3-7). right; mean fluorescence intensity of cytoplasmic region/biological replicate. **b** and **c,** Statistical analysis were performed using pairwise *t*-test with holm adjustment (**P<0.01). **d**, left; Diagram of inhibition of thermogenesis. right; Decreased relative temperature by inhibition of proton leak using BgTEMP-NLS. Relative values in each cell were plotted (DMSO; n = 57 cells, Rot & Ant A; n = 64 cells). **e**, Proton leak in DMSO and Genipin treated BMDM incubated at 28°C and 37°C (n=3 biological replicates/group). **f**, qPCR analysis of gene expression levels for *Aac1*, *Aac2*, *Ucp1*, *Ucp2*, *Ucp3*, *Ucp4*, *Ucp5*, *Slc25a30* in BMDMs incubated at 28°C and 37°C (n = 4-7 biological replicates/ group). **d, e, f,** Statistical analysis were performed using Student’s *t*-test (*P<0.05, **P<0.01). **g**, AAC linked proton leak index (n = 6 biological replicates/ group). Statistical analysis were performed using paired Student’s *t*-test (*P<0.05). In **a**, Data are expressed as means; error bars, s.d. In **b**, **e**, **f** and **g**, box and whisker plots show the the 75th 50th, and 25th percentiles of the data, and minimum and maximum values.

To confirm whether BMDMs actively generate heat via proton leak at 28°C, we employed the ratiometric genetically encoded fluorescent thermometer B-gTEMP (Lu et al., 2022) to measure intracellular temperature. BMDMs were transfected with the plasmid encoding nucleus targeted B-gTEMP and first imaged at 37°C to establish a baseline reference for B-gTEMP fluorescence. Following this, the cells were treated with either Rotenone and Antimycin A or DMSO (control). Rotenone and Antimycin A inhibit proton flow into the cytoplasm by targeting mitochondrial complexes I and III, respectively, thereby suppressing proton leak from the cytoplasm into the mitochondrial matrix. After 4 hours of incubation at 28°C, we recorded B-gTEMP fluorescence and calculated relative temperature by normalizing the 28°C fluorescence with the 37°C reference. BMDMs treated with Rotenone and Antimycin A exhibited a significantly lower relative temperature compared to DMSO-treated cells, indicating proton leak elicited thermogenesis contributes to maintain intracellular temperature (Fig. 2d, sFig. 3a). A similar decrease in intracellular temperature was confirmed using a different thermo-probe (sFig. 3b). In contrast, when cells were incubated at 37°C for 4 hours following the same treatments, no significant difference in relative temperature was observed between the Rotenone/Antimycin A and DMSO groups (sFig. 3c). These results demonstrate that BMDMs generate heat via proton leak specifically under cold conditions (28°C), but not at normothermic conditions (37°C). Although the amount of proton leak was larger at 37°C than that at 28°C on XF analysis (Fig. 2a), this elevated proton leak at physiological temperature may serve a protective role by limiting mitochondrial reactive oxygen species (ROS) production, a mechanism previously implicated in reducing endothelial cell activation and inflammation (Nanayakkara et al., 2019). Importantly, treatment at 37°C with Rotenone and Antimycin A did not significantly reduce intracellular temperature (sFig. 3c), indicating that the observed changes in proton leak are not due to shifts in overall cell temperature, but rather reflect active mitochondrial regulatory processes.

It is known that increased thermogenesis by cold stimulation mainly cause via UCP1, which mediate mitochondrial proton leak, in brown adipocyte and beige adipocyte (reviewed in Bertholet & Kirichok 2021). However, macrophages predominantly express UCP2, rather than UCP1 (Larrouy et al., 1997). To assess the role of UCP2 for proton leak index, we treated BMDMs with the UCP2 inhibitor Genipin. Genipin treatment did not significantly alter proton leak (Fig. 2e), suggesting that UCP2 does not contribute to the increased proton leak index observed at 28°C. Recent studies have shown that ADP/ATP carriers (AACs), which are members of the SLC25 mitochondrial carrier superfamily, can mediate proton leak (Bertholet et al., 2019, Bertholet et al., 2022). Notably, UCP1 and UCP2 are also part of this superfamily. It has been proposed that all SLC25 family members may contribute to proton leak to some extent (Bertholet et al. 2019). To explore the potential involvement of AACs and UCPs in proton leak index under cold conditions, we examined RNA expression levels. At 28°C, expressions of *Aac1* and *Aac2*, which function as ATP/ADP exchangers across the inner mitochondrial membrane, were significantly increased compared to 37°C, whereas UCPs expression did not change (Fig. 2f). AACs become a candidate for keeping thermogenesis via proton leak. Next, to investigate the contribution of AACs to the proton leak index, we inhibited AACs using carboxyatractyloside (CATR). Because inhibition of AACs also blocks ATP/ADP exchange across the mitochondrial inner membrane, CATR treatment reduces ATP-dependent basal respiration in addition to suppressing AAC-mediated proton leak. To isolate AAC-linked proton leak, we subtracted the proton leak measured in CATR-treated BMDMs from that in DMSO-treated controls (Fig. 2g left). The AAC-linked proton leak index was significantly higher at 28°C than at 37°C (Fig. 2g right), indicating that AACs contribute to the maintenance of proton leak under physiological cold conditions through the cold-induced upregulation of AAC1 and 2. Cold stimulation into alternatively activated macrophages has been shown to induce catecholamine production, which subsequently activates thermogenic and β-oxidation genes in brown adipocytes, thereby promoting non-shivering thermogenesis (Nguyen et al. 2011). Sympathetic neuron-associated macrophages also mediate the thermogenesis and *Ucp1* expression by clearance of norepinephrine (Pirzgalska et al 2017). Although indirect contribution of macrophages into the thermogenesis has been already reported, our findings suggest that macrophages, specifically BMDMs, may contribute directly to thermogenesis through sustained proton leak under physiological cold conditions. This highlights a previously underappreciated, cell-autonomous role for macrophages in maintaining thermogenic capacity during cold stress. The existence of thermogenesis pathways that do not rely on neural and hormonal stimulus implies the possibility of local and organ-specific temperature regulation, similar to the specialized heater organs found in certain fishes (Costa and Landeira-Fernandez, 2009).

Physiological cold stimulation reduced mitochondrial membrane potential (Fig. 2c). Previous work by Sanin et al. (2018) demonstrated that mitochondrial membrane potential regulates the expression of Resistin-like molecule alpha (*Retnla*), a key marker of interleukin-4 (IL-4)-activated macrophages. These reports suggest that cold stimulation suppress the *Retnla* expression through decreased mitochondrial membrane potential. In addition, RETNLA has been previously reported to exert antimicrobial effects in the skin (Harris et al. 2019). Thus, cold-induced suppression of RETNLA may have a negative effect of antimicrobial property in mice. To assess how this affects *Retnla* expression, we treated BMDMs with IL-4 and measured *Retnla* levels under both normothermic (37°C) and physiological cold (28°C) conditions. *Retnla* expression was markedly reduced at 28°C compared to 37°C (Fig. 3a, upper panel). RETNLA protein expression was also enhanced at 37°C and was suppressed under cold stimulation (Fig. 3a, lower panel, sFig. 4). To further test whether decreased membrane potential alone could suppress *Retnla* expression, we treated BMDMs at 37°C with uncoupling agents FCCP and valinomycin. These agents, which disrupt mitochondrial membrane potential, similarly inhibited IL-4-induced *Retnla* expression, mimicking the effect of 28°C incubation. These results show that cold-induced suppression of *Retnla* expression occurs through decreased mitochondrial membrane potential. This suppression of *Retnla* expression at 28°C was not related to β-adrenergic receptors (sFig. 5), unlike the thermogenic response of adipocytes. Notably, expression of other IL-4-induced genes *Ccl17* and *Ccl22* was not affected by physiological cold exposure (Fig. 3b), indicating selective regulation. BMDMs isolated from *Stat6* knockout mice did not show increased *Retnla* expression by IL-4. The fold induction of *Retnla* (comparing IL-4-treated versus untreated BMDMs) was similar at 37°C and 28°C. Considering together with similar expression of STAT6 protein expression and phosphorylation level of STAT6 between 37°C and 28°C (Fig. 3c, sFig. 6), it suggests that the IL-4–STAT6 signaling pathway remained equally active at both temperatures. Thus, the observed suppression of *Retnla* at 28°C appears to be due to reduced basal *Retnla* expression prior to IL-4 stimulation, rather than impaired IL-4 signaling itself. A declining in mitochondrial membrane potential is thought to regulate the expression of a subset of genes through potential-dependent mechanisms, such as the transporting DNA-binding proteins from the mitochondria to the nucleus (Sanin et al., 2018). These findings highlight a mitochondria-dependent mechanism of immunometabolic regulation under physiological cold stress.

**Fig. 3.**
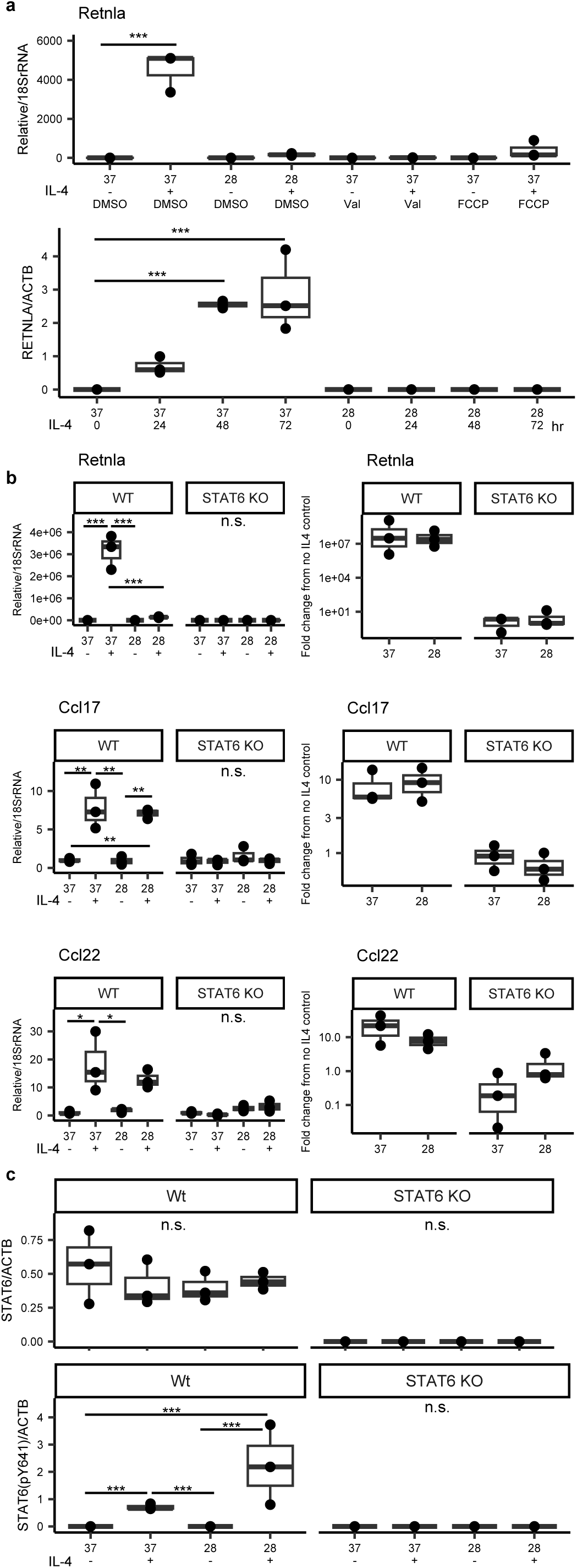
Suppression of *Retnla* expression in BMDMs by cold stimulation. **a, upper panel;** qPCR analysis of gene expression levels for *Retnla* in BMDMs treated DMSO, CATR (6µM), and Valinomycin (50nM). Expression data were corrected by 18S rRNA expression. + means IL-4 treatment, - is not. Multiple comparisons using Dunnett’s test were performed with 37°C, IL-4 -,and DMSO as the reference groups (***P<0.001). **lower panel;** Western blot analysis showing the level of RETNLA in BMDMs incubated at 28°C and 37°C for 0-72 hours after IL-4 treatment. Data were standardized by beta-ACTIN level. Multiple comparisons using Dunnett’s test were performed with 37 and IL-4 at 0 hours as the reference groups (***P<0.001). **b**, qPCR analysis of gene expression levels for *Retnla*, *Ccl17*, and *Ccl22* in BMDMs incubated at 28°C and 37°C (left panel). Right panels indicate fold change from no IL-4 control. **c,** Western blot analysis showing the level of STAT6 and pSTAT6 (pY641) in BMDM incubated at 28°C and 37°C standardized by beta-ACTIN level. **b, c,** Statistical analysis were performed using pairwise *t*-test with holm adjustment (*P<0.05, **P<0.01, ***P<0.001). Box and whisker plots show the 75th 50th, and 25th percentiles of the data, and minimum and maximum values. All plots represent biological replicates (n = 3/group).

To evaluate whether suppression of RETNLA compromises antibacterial activity in vivo, we first tested recombinant RETNLA protein against 23 bacterial strains (Fig. 4a). Among these, only *Acinetobacter baumannii* (ATCC 19606), a Gram-negative and multidrug-resistant pathogen, exhibited susceptibility to RETNLA, whereas no detectable growth inhibition was observed for the other Gram-negative or Gram-positive bacteria tested. Although the molecular mechanism underlying RETNLA-mediated killing of *A. baumannii* remains unclear, this result indicates a species-specific bactericidal activity rather than a broad-spectrum antimicrobial effect. We confirmed that RETNLA exerts a concentration-dependent bactericidal effect against *A. baumannii* (Fig. 4b). RETNLA at 2.5 µM effectively reduced viable bacterial counts, with a detectable lower limit of activity at approximately 0.5 µM (Fig. 4c). As RETNLA is a secreted protein, we next quantified its concentration in the culture supernatant of IL-4-activated BMDMs. After 48 hours of activation at 37°C, RETNLA levels reached ∼0.5 µM, and gradually increased to approximately ∼2 µM by 96 hours, overlapping with the bactericidal concentration range observed in vitro (Fig. 4d, sFig. 7a). Supernatants collected 96 hours after IL-4 stimulation from wild-type BMDMs cultured at 37 °C exhibited significant antibacterial activity against *A. baumannii* than those from *Retnla*-knockout BMDMs (Fig. 4e, sFig. 7b), indicating that RETNLA is a major contributor to macrophage-mediated killing of *A. baumannii* under normothermic conditions. In contrast, supernatants from IL-4-activated BMDMs cultured at 28 °C showed no difference in antibacterial activity between wild-type and *Retnla*-deficient cells, consistent with cold-induced suppression of RETNLA expression. To assess the *in vivo* relevance of RETNLA-mediated antibacterial defense, *A. baumannii* was topically applied to the shaved dorsal skin of mice. No bacteria were recovered following PBS application (Fig. 4f). After 2 days, both RETNLA deficiency and cold exposure significantly attenuated the antibacterial response to *A. baumannii* by two-way ANOVA (mouse genotype x temperature) (Fig. 4g). Skin from *Retnla*-knockout mice tended to harbor higher bacterial burdens than that from wild-type littermates, indicating impaired bacterial clearance (Fig. 4g). Additionally, housing of mice at 16°C markedly suppressed bacterial clearance compared with room temperature housing. These results suggest that RETNLA contributes to antibacterial defense in superficial tissues, while indicating that additional temperature-dependent host factors may also influence bacterial clearance *in vivo*.

**Fig. 4.**
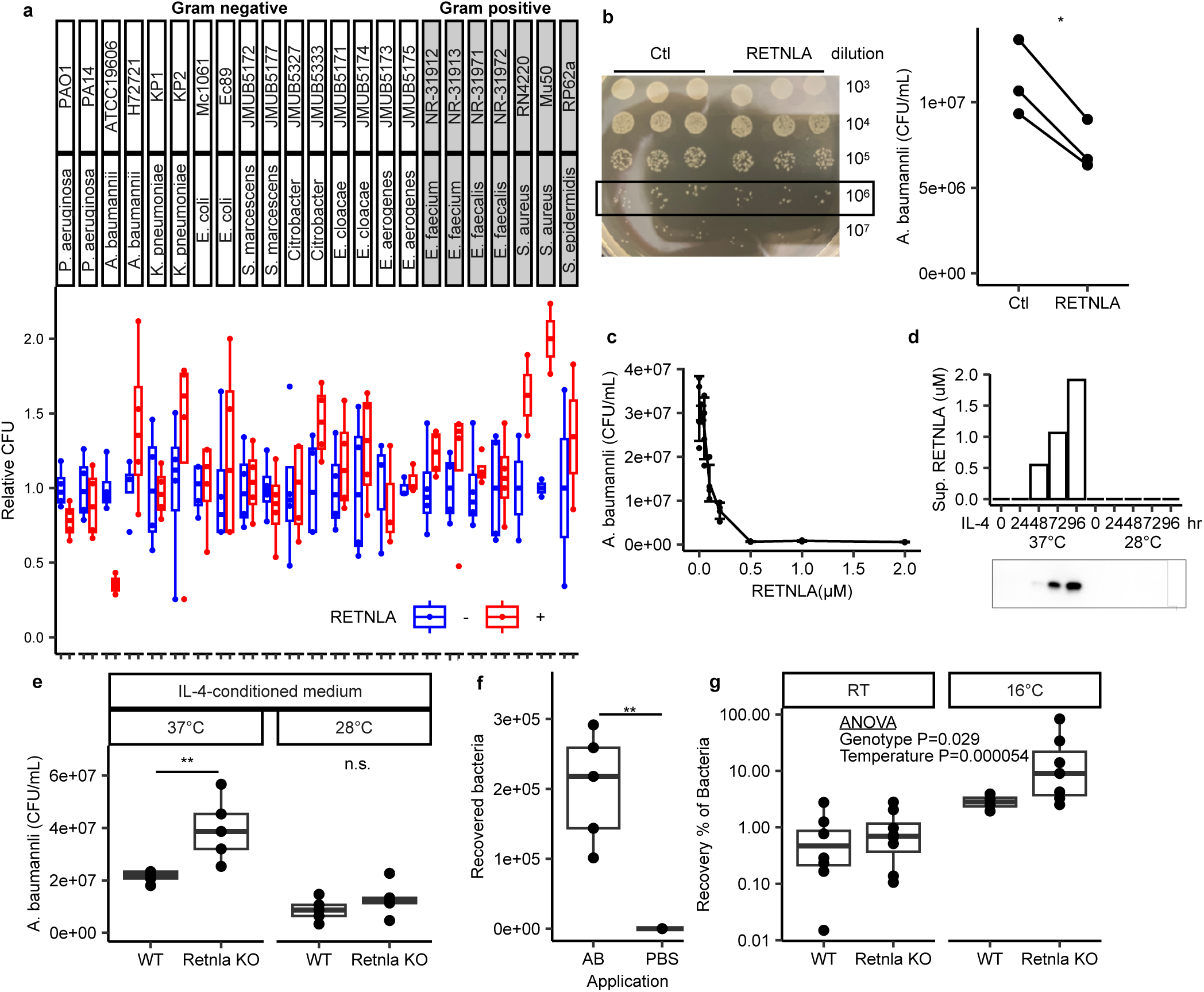
Decreased antibiotics by delation of *Retnla*. **a**, Effect of RETNLA recombinant protein for bacterial strains. Red and blue shows culturing with/ without RETNLA, respectively. Data represents a technical replicate (n=3-4). **b,** Antimicrobial effect of RETNLA for *Acinetobacter baumannii* (ATCC 19606). *A. baumannii* was cultured in the presence of RETNLA (5 µM) or vehicle control (Ctl). Bacterial growth was assessed by serial dilution and plating on LB agar. Colonies at the 10^6^ dilution are counted (boxed area). **Left**; representative image from one experiment. **Right**; points represent experimental means, and connecting lines indicate the same experiment (n=3 independent experimental replicates / group). Statistical analysis was performed using paired Student’s *t*-test. **c**, Lower limit of RETNLA for *A. baumannii* (ATCC 19606). **d**, Western blot analysis showing the level of RETNLA secreted from BMDMs incubated for 0-96 hours after IL-4 treatment (+, not -). **e**, Antimicrobial effect of BMDM-conditioned medium from *Retnla* KO and WT for *A. baumannii* (ATCC 19606). BMDM-conditioned medium was collected after 96 hours of IL-4 treatment from cells incubated at either 28°C or 37°C. *A. baumannii* was incubated with BMDM-conditioned medium for 5 hours. Data represents experimental replicates (n=5/group). Student’s *t*-test was performed (**P<0.01). **f**, Recovery number of Bacteria applied *A. baumannii*. or PBS medium to back skin of B6 mice. Biological replicate n=5/group. Student’s *t*-test was performed (**P<0.01). **g**, Recovery percentage of Bacteria applied to back skin of *Retnla* KO and WT mice after 2 days. Two-way ANOVA were performed to evaluate the effect of Genotype and housing temperature. Room temperature (RT) WT, RT KO, 16°C WT, and 16°C KO mice were n=8, 8, 4, and 7, respectively. In **a**, **e**, **f**, and **g**, box and whisker plots show the the 75th 50th, and 25th percentiles of the data, and minimum and maximum values. In **c**, data are expressed as means; error bars, s.d.

In summary, we demonstrated that BMDMs uniquely respond to physiological cold stress by increasing their proton leak index and suppression of *Retnla* expression. White and beige adipocytes are known to autonomously upregulate thermogenic genes in response to cold exposure (Ye et al, 2013). Even in ectotherms, cells autonomously control intracellular temperature by the enhancement of F_1_ F_0_-ATPase-dependent mitochondrial respiration (Murakami et al, 2022). Similarly, BMDMs exhibited a cell-autonomous mechanism of heat production through AAC-linked proton leak. Cold exposure has been shown to impair immune response in nasal cavity (Foxman et al, 2015) and damage the specific organ (Guo et al, 2022). In this context, cell-autonomously thermogenesis of macrophages may serve as a protective mechanism, buffering the functional decline of peripheral tissues and organ under physiological cold conditions. Given the cold-induced suppression of RETNLA-mediated antibacterial activity, this thermogenic mechanism may be critical for sustaining immune function and cellular homeostasis in variable temperature environments. Conversely, macrophages in peripheral and superficial tissues might also be prepared for heat-induced RETNLA activation, since inflammation caused by bacterial infection or injury generally increases local temperature. While non-shivering thermogenesis is traditionally attributed to UCP1-dependent mechanisms in adipocytes, UCP1-independent thermogenic pathways involving SERCA and AAC have also been proposed (Roesler & Kazak, 2020). AAC-linked proton leak has previously been reported in mitoplasts from smooth muscle, heart, kidney, and liver (Bertholet et al, 2019). Our findings extend this concept to macrophages, revealing a novel role for immune cells in peripheral and localized thermogenesis. Thus, peripheral macrophage in mice may function to help maintain local temperature, thereby buffering of the cold-induced reduction of *Retnla* expression and antimicrobial activity under cold conditions.

## Materials & Methods

### Ethics statement

All animal experiments were carried out in a humane manner. The Institutional Animal Experiment Committees of Jichi Medical University and The University of Tokyo approved this study. This study was conducted in accordance with the Institutional Regulations for Animal Experiments and Fundamental Guidelines for Proper Conduct of Animal Experiments and Related Activities in Academic Research Institutions under the jurisdiction of the MEXT of Japan.

### Animals

Six-week-old male C57BL/6J mice were purchased from CLEA Japan and were housed until the experimental day. B6N.129S6-Retnla^tm1Tliu^/J *(Retnla* knockout *mice,* IMSR_JAX:029976, Liu et al 2014) and *Stat6^tm1Gru^* (*Stat6* knockout mice, IMSR_JAX:002828, Kalpan et al 1996) were originally obtained from The Jackson Laboratory. Heterozygous *Retnla* knockout mice, *Stat6* knockout mice, and GO-ATeam-expressing mice (AVID mouse, Ohnishi et. al., 2025) were inbred. All mice were housed under a 12-hour light/dark cycle in a temperature-controlled room (22°C±2°C), with ad libitum access to food and water.

### Cells

#### Harvest of Mouse Bone Marrow-Derived Macrophages (BMDMs)

Bone marrow cells were isolated from the femurs and tibias of 6–7-week-old C57BL/6J mice on day 0. Cells were seeded in RPMI-1640 medium supplemented with 10% heat-inactivated fetal bovine serum (FBS), 1% penicillin-streptomycin, and 30% conditioned medium (supernatant from M-CSF-expressing L929 cells). The culture medium was refreshed every two days. On day 6, the medium was replaced with RPMI1640 supplemented with 10% heat-inactivated FBS, 1% penicillin-streptomycin, and 20 ng/mL recombinant M-CSF (BioLegend). Mature BMDMs were used for experiments from day 6 onward. IL-4 (RSD) stimulation was performed at final 10µg/mL.

### Other Cell Lines and Differentiation Protocols

Epithelial like cell NCTC1469 (JCRB9075) and preadipocytes 3T3-L1 (IFO50416) was cultured in DMEM supplemented with 10% heat-inactivated FBS, 1% penicillin-streptomycin. Differentiation of preadipocytes 3T3-L1 into adipocyte was induced in the DMEM medium containing 1 mM IBMX (Sigma, I5879), 1µM dexamethasone (Wako 047-18863), 100nM insulin (Sigma, I6634) for 8 days.

Preadipocytes T37i (Sigma, SCC250) was cultured in DMEM/F12 supplemented with 10% heat-inactivated FBS, 1% penicillin-streptomycin. Differentiation into adipocyte was induced in the DMEM/F12 medium containing 8nM 3,3’,5-Triiodo-L-thyronine sodium salt (Wako), 80nM insulin for 8 days. Pre-keratinocyte PHK16-0b (JCRB0141) was cultured in KGM-Gold Keratinocyte Growth Medium BulletKit (LONZA, 00192060). Differentiation into keratinocyte was induced for 5 or 6 days in the KGM-Gold Keratinocyte Growth Medium containing 1.8 mM CaCl_2_.

### Quantitative PCR

Total RNA was extracted using the FastGene RNA Basic Kit (NIPPON Genetics) in combination with QIAshredder columns (QIAGEN), following the manufacturers’ protocols. Complementary DNA (cDNA) was synthesized from total RNA using the RevaTraAce qPCR RT Master Mix (TOYOBO), according to the manufacturer’s instructions. Real-time qPCR was performed using Thunderbird SYBR qPCR Mix (TOYOBO) and Light cycler 480 (Roche) with specific primers for each gene (Table 1). Standard curves generated from serial dilutions of pooled cDNA were used to determine primer amplification efficiency. Relative gene expression levels were calculated using the Abs Quant/Second Derivative Max method. 18S rRNA was used as the internal control for normalization.

**Table 1.**
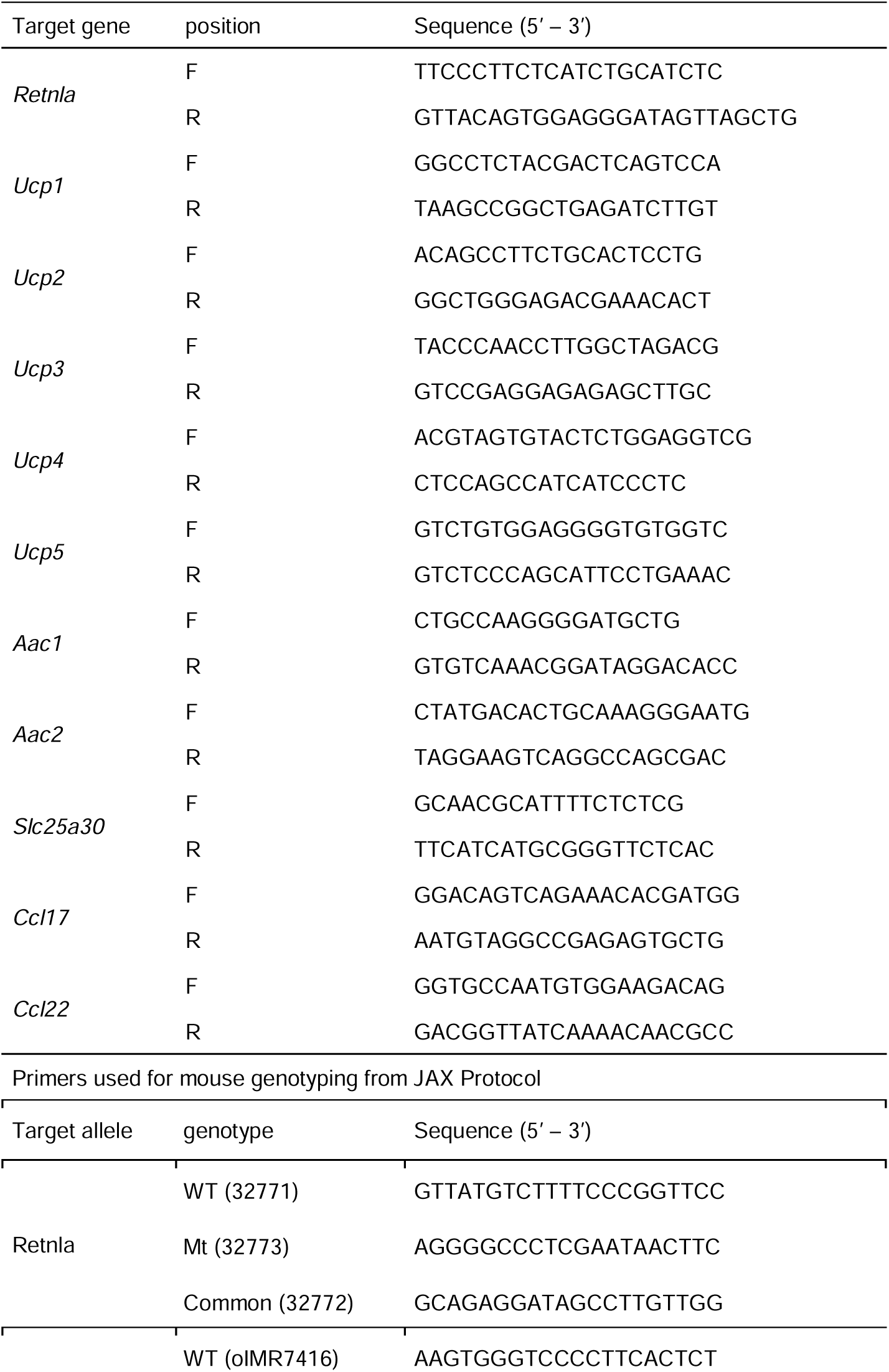
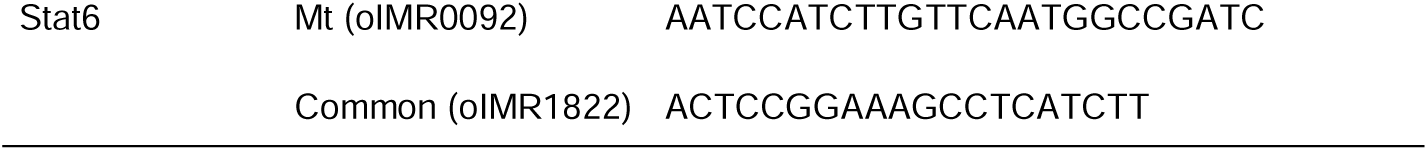
Primers used for qPCR.

### Western blotting

BMDMs were lysed in RIPA buffer containing 0.5% SDS supplemented with protease (Roche, 04693116001) and phosphatase (nacalai, 07575-51) inhibitors. Lysis was performed on ice for 30 minutes, followed by centrifugation at 15,000 × *g* for 15 minutes at 4°C. Protein concentration in the supernatant was measured using the Pierce™ Rapid Gold BCA Protein Assay Kit (Thermo Scientific, A55860).

For immunoblotting, 1 µg of total protein per sample was separated by SDS-PAGE and transferred to a PVDF membrane (Cytiva, GE10600021). The following primary antibodies were used: the STAT6 (1:1000 dilution,Cell Signaling Technology #9362S), Retnla (1:1000 dilution, Abcam ab39626), internal control ATCB (MBL, M177-3). Detection was performed using ECL Prime Western Blotting Detection Reagent (Cytiva, RPN2232). Band intensities were quantified using FIJI/ImageJ, according to standard protocols.

### Metabolome analysis

Matured BMDMs were incubated at 28°C for 4 hours or 24 hours and 37°C for 24 hours from the day 6. At day 7, cells were washed by PBS twice and immediately frozen by liquid nitrogen. From frozen BMDMs samples, metabolites were prepared as following published protocols (Maeda et al, 2023). Frozen BMDMs were sonicated in ice-cold methanol containing internal standard compounds. Methionine sulfone (L-Met) and 2-morpholinoethanesulfonic acid (MES) were used as internal standards for cationic and anionic metabolites, respectively. An equal volume of chloroform and 0.4 times the volume of ultrapure water (LC/MS grade, Wako Pure Chemical, Tokyo, Japan) were added. After centrifugation, the aqueous phase was collected and filtered through an ultrafiltration tube (Ultrafree MC-PLHCC, Human Metabolome Technologies, Tsuruoka, Japan). The filtrate was concentrated using a vacuum concentrator (SpeedVac; Thermo Fisher Scientific, Waltham, MA, USA), dissolved in 50 µL ultrapure water, and subjected to ion chromatography–high-resolution mass spectrometry (IC-HR-MS) analysis. Recovery rates (%) of these standards were used to correct for the loss of endogenous metabolites during sample preparation.

#### Ion chromatography-high resolution mass spectrometry (IC-HR-MS) metabolome analysis

Metabolites were detected using an Orbitrap-type MS instrument (Q-Exactive focus; Thermo Fisher Scientific) coupled to a high-performance ion chromatography system (ICS-5000□+□, Thermo Fisher Scientific). The IC instrument was equipped with an anion electrolytic suppressor (Dionex AERS 500; Thermo Fisher Scientific) to convert the potassium hydroxide gradient into pure water prior to MS detection. Separation was performed using a Dionex IonPac AS11-HC column (4 µm particle size, Thermo Fisher Scientific). The IC flow rate was 0.25□mL/min, supplemented post-column with 0.18□mL/min makeup flow of MeOH. The potassium hydroxide gradient was programmed as follows: 1 mM to 100 mM (0–40 min), held at 100 mM (40–50 min), then returned to 1 mM (50.1–60 min). Column temperature was maintained at 30°C. The mass spectrometer was operated in the ESI-negative mode for all detections. A full mass scan (m/z 70–900) was performed at a resolution of 70,000. The automatic gain control target was set at 3 × 10^6^□ions, with a maximum injection time of 100 □ms. The source ionization parameters were: spray voltage of 3□kV, transfer temperature, 320□°C; S-Lens level, 50; heater temperature 300□°C; sheath gas 36; and aux gas, 10.

#### Liquid chromatography-tandem mass spectrometry (LC-MS/MS) for amino acid measurement

Cationic metabolite concentrations were determined by LC-MS/MS. A triple-quadrupole mass spectrometer equipped with an electrospray ionization (ESI) source (LCMS-8060, Shimadzu Corporation, Kyoto, Japan) was operated in both positive and negative ion modes using multiple reaction monitoring (MRM). Separation was achieved on a Discovery HS F5-3 column (2.1□mm I.D. × 150□mm□L, 3 μm particle size; Sigma-Aldrich, St. Louis, MO, USA) with a gradient elution of mobile phase A (0.1% formate) and mobile phase B (acetonitrile containing 0.1% formate). The elution gradient was: 100:0 (0–5 min), 75:25 (5–11 min), 65:35 (11–15 min), 5:95 (15–20 min), and 100:0 (20–25 min). Flow rate was maintained at 0.25 mL/min, and column temperature at 40°C.

#### Quantitative and qualitative data analyses

Metabolites registered in our in-house compound library were identified by comparison with authentic standards, confirming retention time and mass accuracy. Chromatographic peak integration and compounds validation were performed for IC-HR-MS and LC-MS/MS, respectively, using Trace Finder software (ver. 4.1, Thermo Fisher Scientific) and Lab Solutions software (ver. 5.113, Shimadzu). For IC-HR-MS, identification parameters were: molecular ion intensity threshold 10000, signal-to-noise ratio ≥ 5, and mass tolerance ≤ 5 ppm. Isotope pattern analysis using a 90% fit threshold, 30% allowable relative intensity deviation, and 5 ppm mass deviation were also performed to ensure that the relative intensities of the M□+□1 and/or M□+□2 isotope peaks for each compound were consistent with the theoretical relative intensities. For LC-MS/MS analysis, chromatographic peaks were integrated using compound-specific SRM channels and manually reviewed. Confirmatory SRM transitions were used when available. Quantitative values were normalized to internal standards (MES for IC-HR-MS and L-Met for LC–MS/MS) to correct for recovery.

#### Multivariate statistical analysis

Supervised partial least squares discriminant analysis (PLS-DA) was conducted using MetaboAnalyst (v4.0) (Chong et. al. 2018). Data were normalized by median adjustment to correct systematic variation across samples and autoscaled to standardize variable ranges.

### B-gTEMP and GO-ATeam

A synthetic cDNA encoding nucleus targeted B-gTEMP was designed by flanking nuclear localization signal sequence (APKKKRKV) to the C-terminus of the B-gTEMP sequence. The synthetic cDNA was PCR-amplified and cloned into pcDNA3.1 vector backbone to construct pcDNA3.1/ B-gTEMP-NLS (Thermo Fisher Scientific, Waltham, MA, USA).

On day 6 of BMDMs, 5 µg pcDNA3.1/B-gTEMP-NLS plasmids were transfected into BMDMs (> 1.0 x 10^6^) with Mouse macrophage Nucleofector Kit (VPA-1009, Lonza) using Nucleofector® 2b (Lonza, program Y-001). Transfected BMDMs were seeded into 3.5 cm dish coated with collagen (Matsunami, D11134H). After 1-2 days, BMDMs were observed by confocal microscopy (Leica, SP8) using oil-immersion objective. B-gTEMP transfected BMDMs were excited with 488 nm laser at the minimal output power of 0.5%. Fluorescence of mNeonGreen (mNG) and tdTomato (tdT) was collected through emission filters 500/520 and 570/590, respectively. The fluorescence intensity was quantified from manually set a region of interest (ROI)/cell using FIJI (ImageJ) (National Institutes of Health, Bethesda, MD, USA) software (Schindelin et al, 2012). On day 6, ATP sensor GO-ATeam expressing mice (Ohnishi et. al., 2025) derived BMDMs were excited with 488 nm laser at the minimal output power of 0.5 %. Fluorescence of mGFP and mKO was collected through emission filters 500/520 and 550/570, respectively. Fluorescence intensity was quantified in FIJI from automatically defined ROI using the default threshold setting. The FRET ratio (mKO/mGFP) was calculated as an ATP indicator.

### Inhibition of cellular thermogenesis

B-gTEMP transfected BMDMs were observed at medium temperature 37°C and 5% CO_2_ controlled with heating cooling chamber (STXFC-KRiX-SET, Tokai Hit). Then Rotenone (4µM) & Antimycin A (1µM) were applied to the dish. The BMDMs were incubated at 28°C and 5% CO_2_ on stage for 4 hours and then were observed. Effect of Rotenone & Antimycin A for thermogenesis was evaluated with Fluorescence ratio (mNG/tdT) at 28°C corrected by Fluorescence ratio (mNG/tdT) at 37°C.

### Oxygen Consumption Rate (OCR) analysis using XF24

Bone marrow cells were seeded by 8 x 10^5^ cells /well in XF24 plate (Agilent) on day 0. The culture medium was changed as described above. On day 6, cell was incubated at either 28°C, 31°C or 37°C with 5% CO_2_ for 24 hours. On day 7, the medium was replaced to Seahorse XF RPMI assay medium supplemented with 20 ng/mL recombinant M-CSF.

For other cell types, NCTC1469, differentiated 3T3-L1, differentiated T37i were seeded by 2.5 x 10^4^ cells per well in XF24 plate. Keratinocyte were differentiated in XF24 microplate for 5 days. Then, the cells were incubated at either 28°C or 37°C for 24 hours in 5% CO_2_. Next day, medium was changed to XF DMEM assay medium. OCR measurements were conducted using the extracellular flux analyzer (Seahorse XFe24 Analyzer, Agilent) at 28°C, 31°C or 37°C. Oligomycin A (final 2.5 μM), Carbonyl cyanide 4-(trifluoromethoxy) phenylhydrazone (FCCP, final 6 μM), and Antimycin A (1 μM) plus Rotenone (Rot, 4 μM) were added in three sequential injections. Median value among technical replicate wells was used for analysis. Mean value in each injection session was used to calculate the following mitochondrial parameters:

Basal respiration = Basal OCR − OCR after antimycin A/rotenone.

ATP-linked respiration = Basal OCR − OCR in oligomycin session.

Proton leak = OCR in oligomycin session − OCR after antimycin A/rotenone.

Maximal respiration = OCR in FCCP session − OCR after antimycin A/rotenone.

Spare respiratory capacity = OCR in FCCP session − Basal OCR.

Proton leak index = Proton leak / Spare respiratory capacity.

To estimate the AAC linked Proton Leak index, cells were treated with the adenine nucleotide translocase (AAC) inhibitor carboxyatractyloside (CATR, final 50 μM, (under CATR treatment), Fig. 2g).

**Non-AAC-linked proton leak** = OCR in oligomycin session (under CATR treatment) − OCR after antimycin A/rotenone (under CATR treatment).

**AAC-linked proton leak** = Proton leak (DMSO control) − Non-AAC-linked proton leak (CATR).

**AAC-linked proton leak index** = AAC-linked proton leak / Spare respiratory capacity (DMSO).

To estimate UCP2 contribution of proton leak, cells were pre-treated with Genepin (final 50µM). Proton leak was calculated under Genepin treatment.

### Membrane potential measurement of Cells

Bone marrow cells and PHK-0b pre-keratinocytes were seeded into CellCarrier Ultra 96-well plates (PerkinElmer) at densities of 1×10^5^ and 2×10^4^ cells per well, respectively, and cultured under differentiation conditions as described. NCTC1469, differentiated 3T3-L1, and differentiated T37i were seeded at 2×10^4^, 2.5×10^4^, and 2.5×10^4^ cells per well, respectively. All cells were incubated with MT-1 (1:1000 dilution, Dojindo) for 30 minutes at 37°C in 5% CO_2_. After incubation, the medium was replaced to each culture medium, and the cells were incubated for additional 24 hours at 37°C, 31°C, or 28°C in 5% CO_2_. Following temperature treatment, cells were fixed with 4% paraformaldehyde and stained with Hoechst 33342 to visualize nuclei. Imaging was performed using Operetta CLS high-content imaging system (PerkinElmer). Nucleus were identified by Hoechst 33342 fluorescence with algorithms B of Operetta software. Cytoplasm was defined by MT-1 fluorescence surrounding nucleus with algorithms A of Operetta. Cytoplasmic area was defined by subtraction between Cytoplasm and Nucleus. Mean intensity of MT-1 fluorescence within cytoplasmic area was obtained as an indicator of relative mitochondrial membrane potential.

### Bacteria Killing assay in *vitro*

Each bacterial strain (Fig 4a) was cultured in either LB or TBS medium for about 8 hours. Cultures were then adjusted to a concentration of 3 × 10□ CFU/mL, and 2.5 µM recombinant RETNLA (PEPROTECH) was added to the medium. After incubation, the antibacterial effect of RETNLA on each strain was assessed by serial dilution and plating on LB or TBS agar plates. Colony-forming units (CFUs) were counted to quantify bacterial growth. Conditioned medium of BMDMs after treatment of IL-4 for 96 hours was used in bacteria killing assay.

### Bacteria Killing assay in *vivo*

Six- to 10-week-old mice were pre-housed at either RT or 16°C for 1 week. After acclimation, dorsal back fur was removed from C57BL6 *Retnla* Knockout and wildtype litter mate mice under anesthesia using shaver followed by depilatory cream (Nair Hair Remover Lotion). Anesthesia was induced with a combination of medetomidine, midazolam, and butorphanol, and reversed with atipamezole. On the following day, the dorsal skin was superficially abraded in a vertical pattern using an 18G needle (TERUMO). Subsequently, 20 µL of *Acinetobacter baumannii* (ATCC19606) in logarithmic growth phase was applied to the abraded area. After 48 hours, mice were euthanized, and the infected dorsal skin was excised and homogenized in sterile PBS. CFUs of bacteria were assessed by serial dilution and plating on LB agar.

### Statistical analysis

All statistical analyses were performed in R (R Core team 2021). Depending on the experimental design, unpaired or paired Student’s *t*-test, two-way ANOVA, Holm-adjusted multiple comparisons or Dunnett’s test were applied as indicated in the figure legends. Sample sizes (n) reported in figure legends refer to the number of replicates shown. For cell imaging experiments, n indicates the number of cell regions of interest analyzed from each experiment. Data was assumed to follow normal distribution, although this was not formally tested. Statistical significance was set at P < 0.05.

## Supporting information

sFig

Sorce Data

Material list

## Acknowledgement

We thank all members of the Division of Bioconvergence for their helpful discussions and the care of mice used in this study. This work was supported by JSPS (Japan Society for the Promotion of Science) KAKENHI (Grant-in-Aid for Scientific Research C), grant number 21K06858 (H.S.), 24K10096 (H.S.), and 25K11306 (T.I.) and AMED (Japan Agency for Medical Research and Development), grant number JP23gm6510023 (N.T.). We used AI-assistant language editing tool (ChatGPT, OpenAI) to improve the readability and grammar of manuscript. All AI-edited text was reviewed and verified by the authors.

## Author contributions

N.T. supervised this study; H.S., T.I., K.K., L.C., and N.T. designed the experiments; H.S., K.M., K.K., Y.S., and T.K. performed the experiments. H.S., T.I., K.M., K.K., Y.S., and N.T. analyzed and interpreted the data; M.Y. provided the GO-ATeam mouse; H.S. wrote manuscript; All authors (H.S., T.I., K.M., K.K., Y.S. I.M., T.K., L.C., and N.T.) made manuscript revisions.

