## Supplementary material for "Cell-autonomous thermogenesis of macrophage alters its antibacterial function": sFig

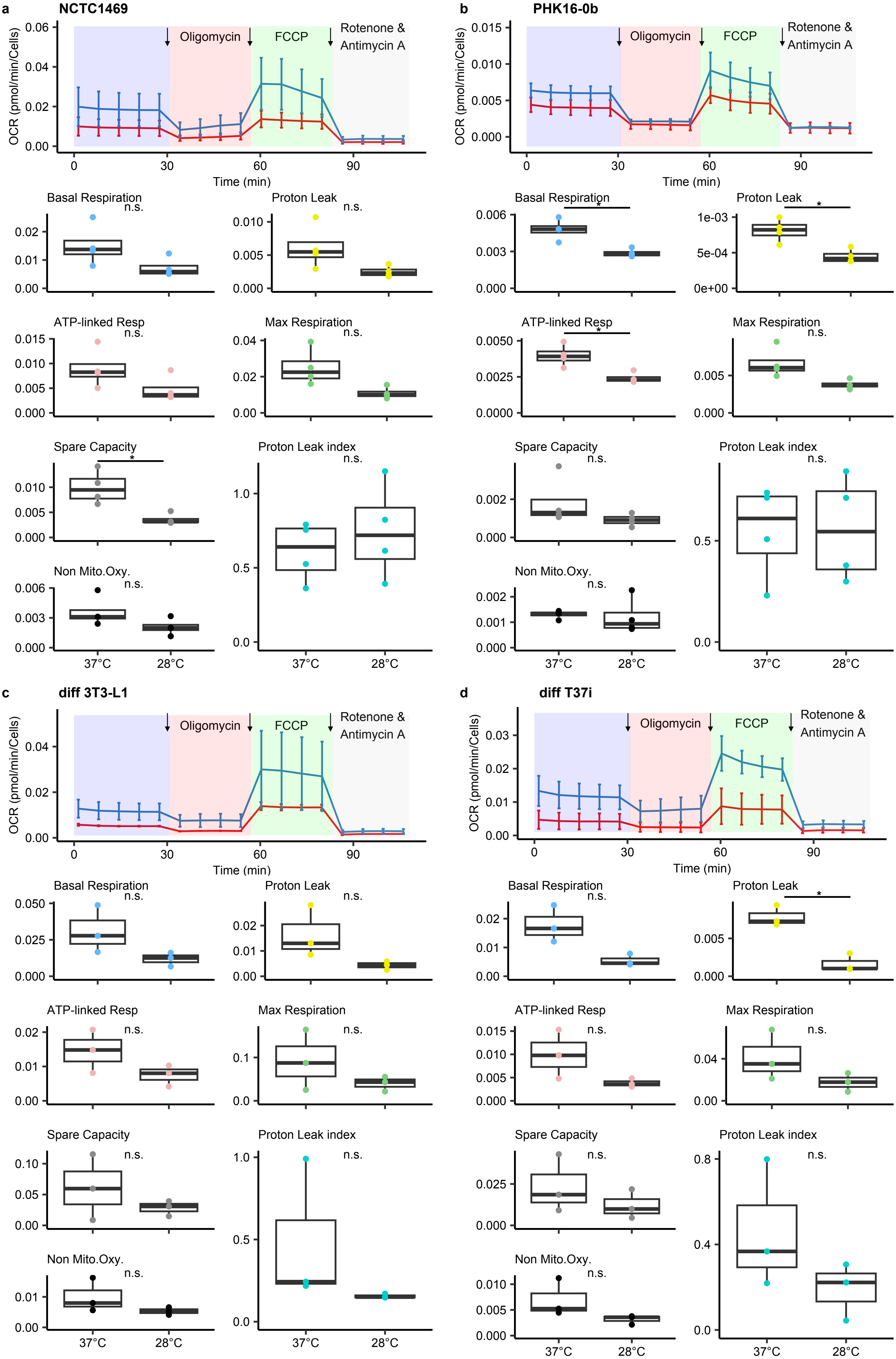
a
NCTC1469
b
PHK16-0b
c
diff 3T3-L1
d
diff T37i
Oligomycin
FCCP
Rotenone &
Antimycin A
OCR (pmol/min/Cells)
Time (min)
Basal Respiration
Proton Leak
ATP-linked Resp
Max Respiration
Spare Capacity
Proton Leak index
Non Mito.Oxy.
n.s.
*
37°C
28°C

sFig. 1

sFig. 1 Oxygen consumption rate (OCR) of mitochondria.
**a**, NCTC1469, epithelial cells. **b**, PHK16-0b, keratinocytes. **c**, differentiated 3T3-L1, white adipocytes, **d**, differentiated T37i, brown adipocytes.
**Upper panel**; OCR, n=3-4 independent experimental replicates/group). Data are expressed as means; error bars, s.d.
**Lower panels**; Quantification plot from each OCR incubated at 28°C and 37°C. Statistical analysis were performed using Student's *t*-test (*P<0.05). Box and whisker plots show the the 75th 50th, and 25th percentiles of the data, and minimum and maximum values.

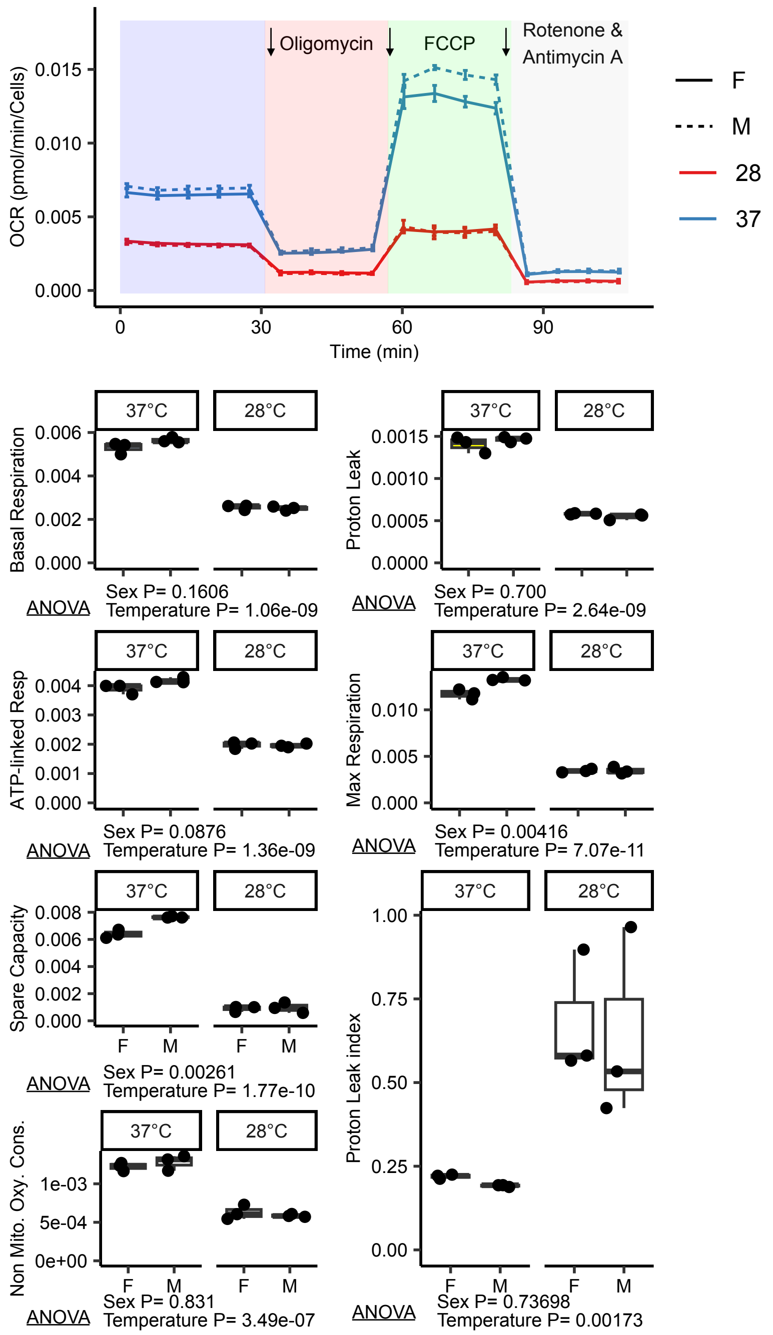


sFig. 2 Sex differences of Oxygen consumption rate (OCR) parameters for cold stimulation. Quantification plot from each OCR of BMDM incubated at 28°C and 37°C (n=3 biological replicates/ group). Two-way analysis of variance (ANOVA) with factors Sex and Temperature was performed. Box and whisker plots show the the 75th 50th, and 25th percentiles of the data, and minimum and maximum values.

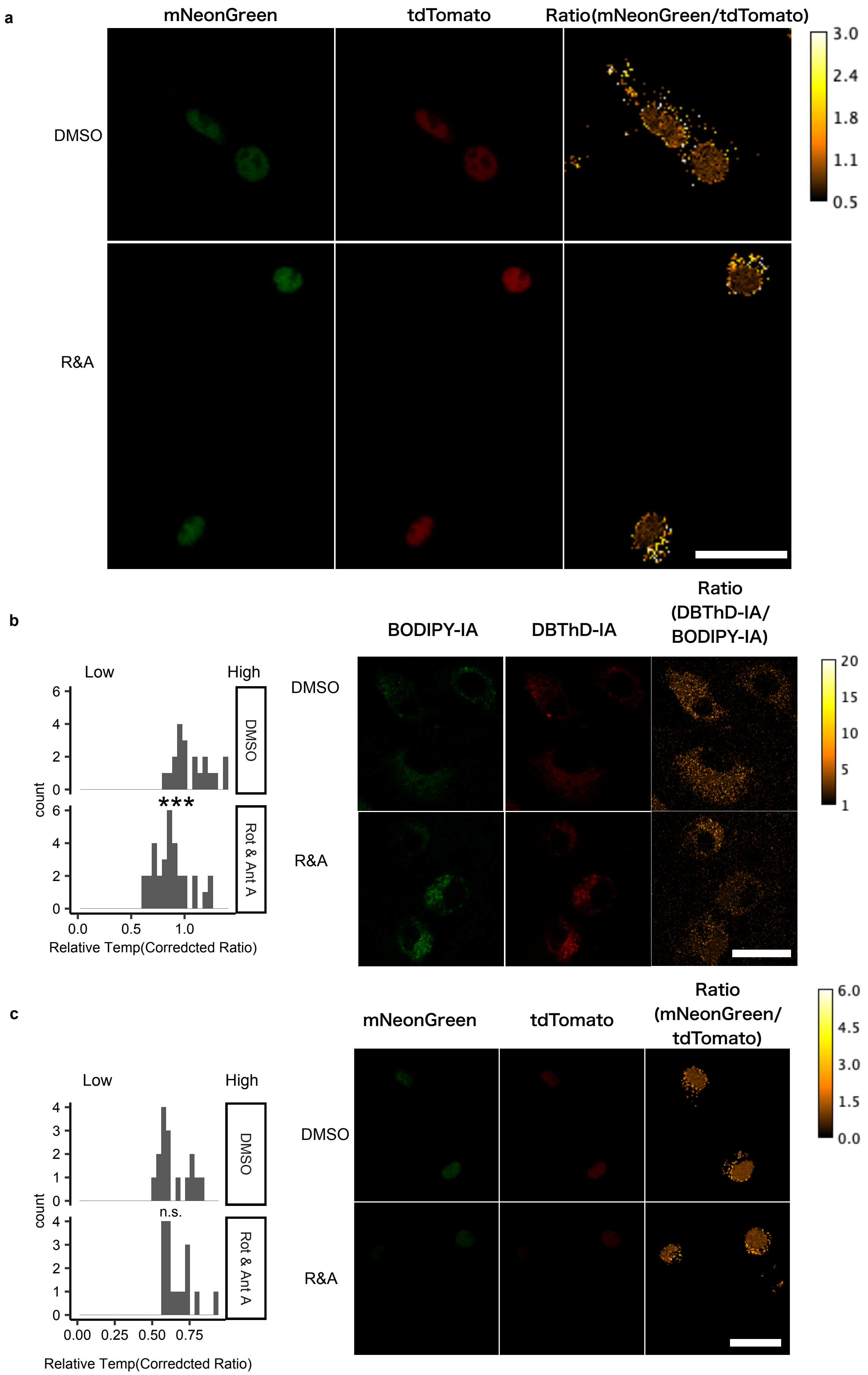


sFig. 3 Decreased relative temperature by inhibition of proton leak using Thermo-probe.
**a**, Representative image of relative temperature after inhibition of proton leak at 28°C using BgTEMP-NLS (Fig. 2d). Bar = 20μm. **b**, Decreased relative temperature by inhibition of proton leak using ThermoProbe® and representative image. Relative values in each cell were plotted (DMSO; n = 20 cells, Rot & Ant A; n = 32 cells). Statistical analysis were performed using Student's *t*-test (***P<0.001). Bar = 20μm.
**c**, Relative temperature after inhibition of proton leak at 37°C using BgTEMP-NLS and representative image. Relative values in each cell were plotted (DMSO; n = 16 cells, Rot & Ant A; n = 16 cells). Statistical analysis were performed using Student's *t*-test. Bar = 20μm.

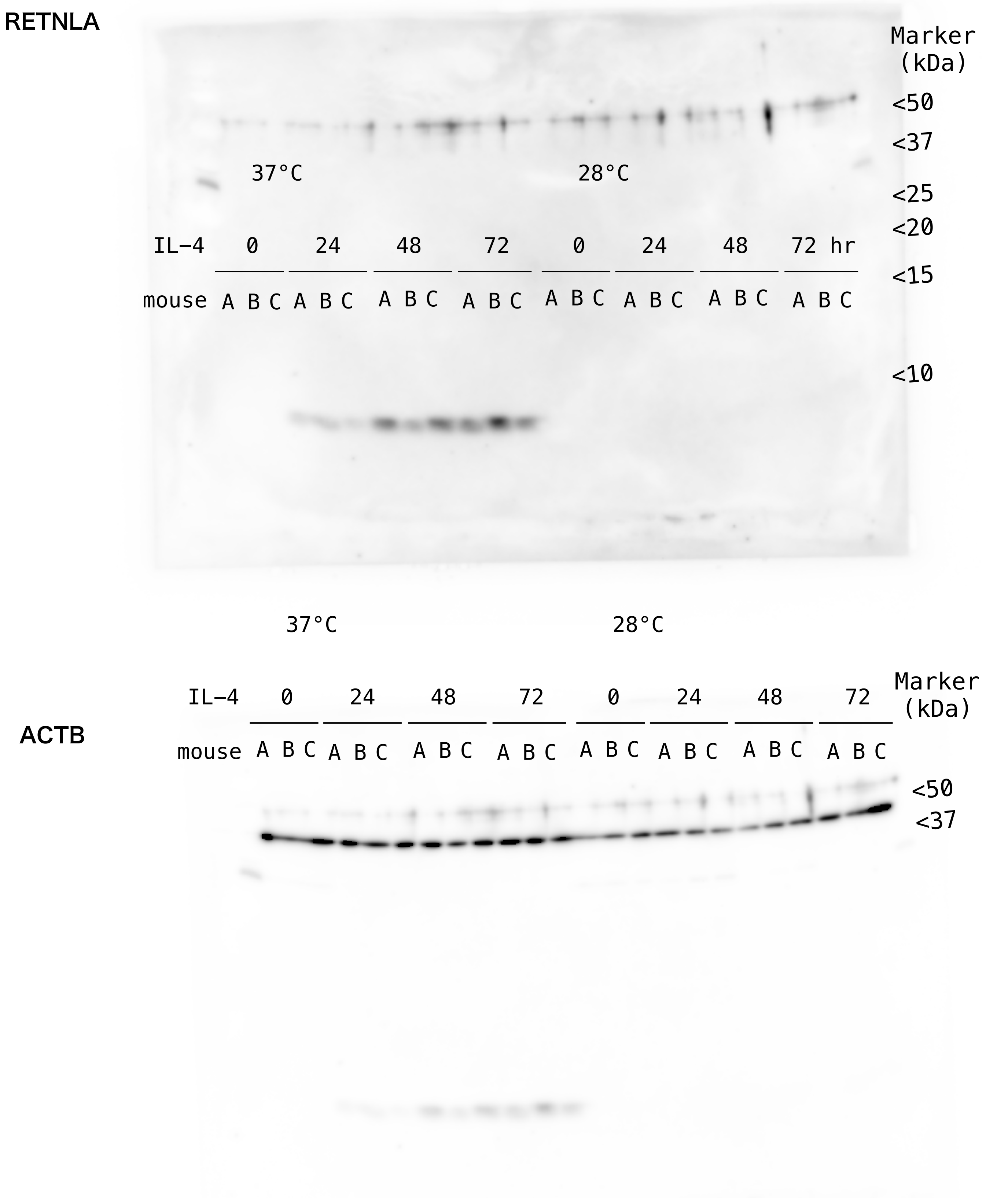


sFig. 4 Western blot of RETNLA (Fig. 3a Lower panel). Upper panel shows the level of RETNLA in BMDM incubated at 28°C and 37°C for 0-72 hours after IL-4 treatment. Lower panel shows beta-ACTIN level as loading control. A,B, and C indicate biological replicates.

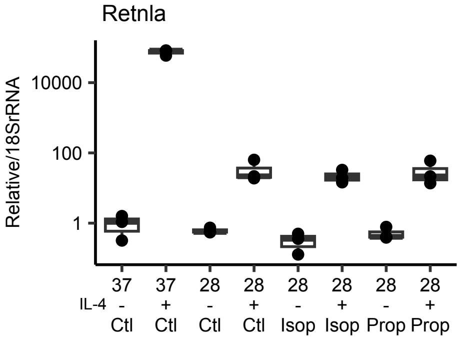


sFig. 5 Adrenergic receptor signaling does not affect *Retnla* expression at 28°C. The beta-adrenergic antagonist propranolol (Prop, 1μM) and beta-adrenergic agonist isoproterenol (Isop, 1μM) was added 1 hour before IL-4 stimulation. + means IL-4 treatment, - is not. Neither treatment altered *Retnla* expression at 28 °C. Box and whisker plots show the 75th 50th, and 25th percentiles of the data, and minimum and maximum values. Plots represent biological replicates (n = 3/group).

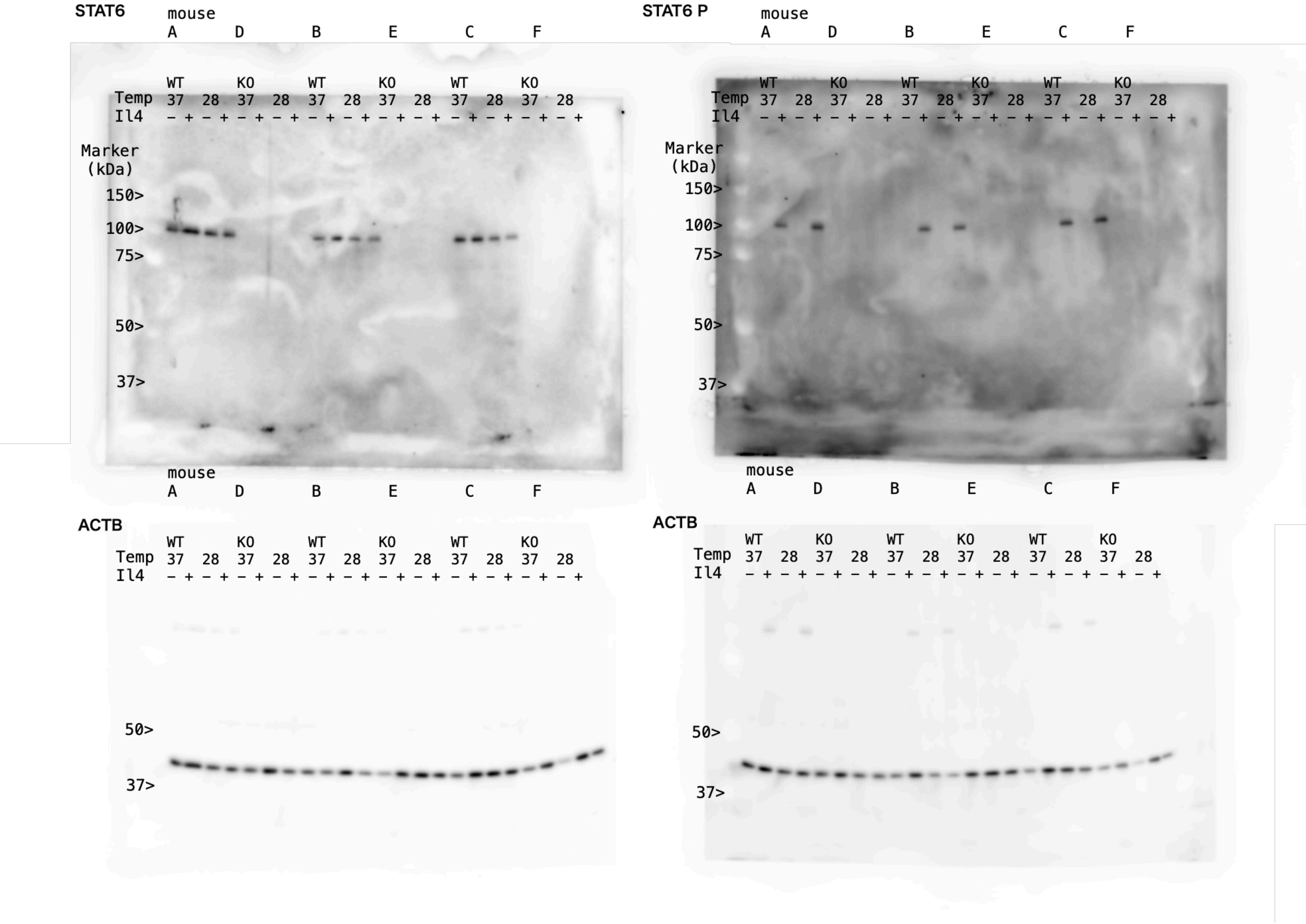


sFig. 6 Western blot of STAT6 and phosphorylated STAT6 (Fig. 3c). Upper panel shows the level of STAT6 (Left) and phosphorylated STAT6 (Right) in BMDM incubated at 28°C and 37°C with IL-4 treatment (+ or -). Lower panel shows beta-ACTIN level as loading control. A, B, C, D, E, and F indicates biological replicates.

**a**

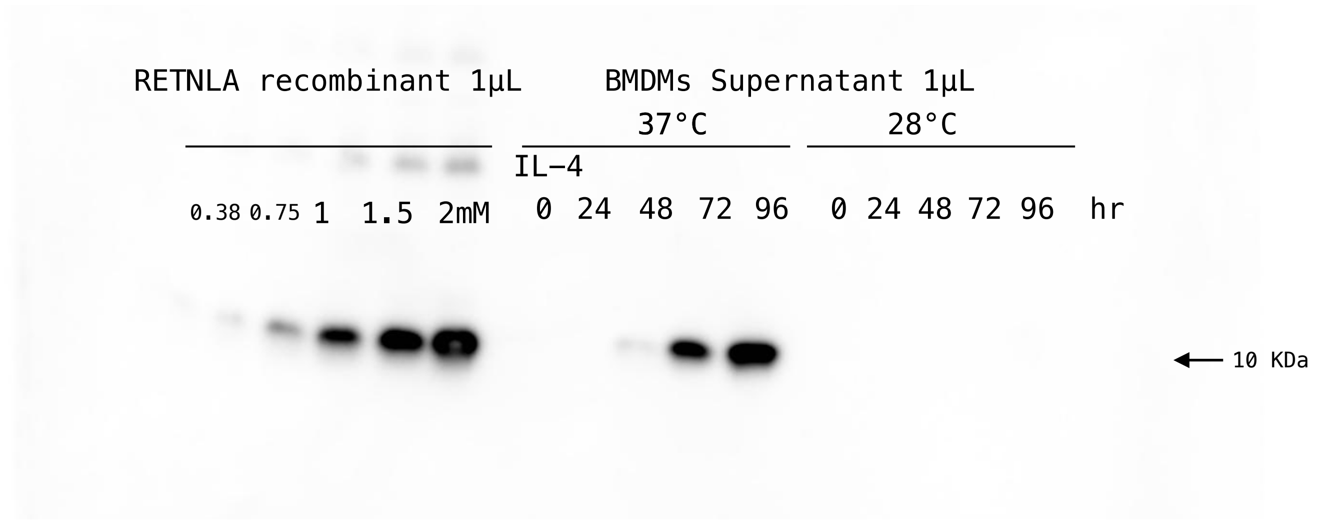


**b**

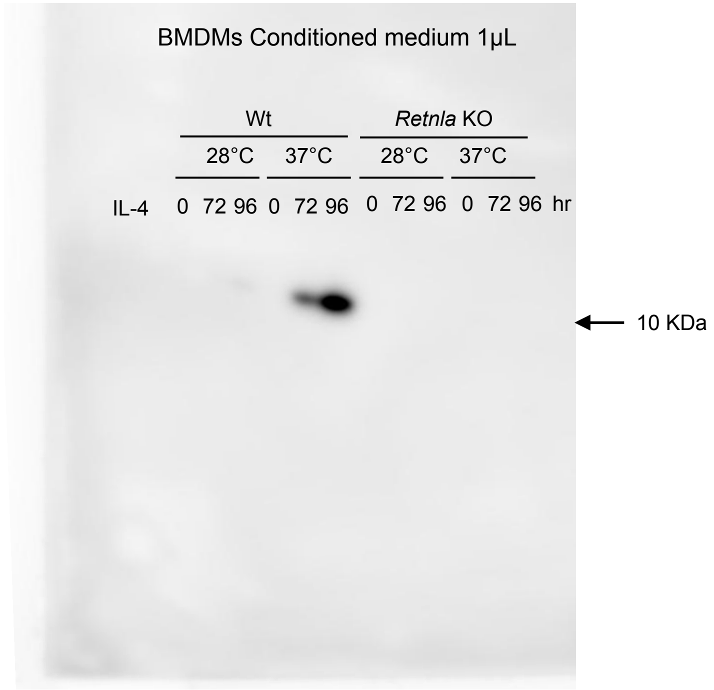


sFig. 7 Western blot of RETNLA.

**a** Conditioned medium of BMDMs (Fig. 4d) from C57BL6/J mice at 37°C and 28°C after IL-4 stimulation.

**b** Conditioned medium of BMDMs (Fig. 4e) from *Retnla* KO and wildtype (Wt) mice at 37°C and 28°C after IL-4 stimulation.
